# Unveiling the diversity, transmission, and zoonotic potential of microbes in true insectivores

**DOI:** 10.1101/2024.12.04.626791

**Authors:** Hongfeng Li, Zheng Y.X. Huang, Jie Lan, Li Hu, Xuemin Wei, Yuhao Wang, Xiujun Li, Yang Li, Daniel J. Becker, Fuwen Wei, Yifei Xu

## Abstract

The Eulipotyphla (true insectivores) is the third largest mammalian order, comprising over 500 species, and could be an important source of human infectious diseases. However, relatively little is known about the diversity of microbes in insectivores and the contribution of insectivores to virus transmission more specifically among wild hosts. In this study, we compiled a comprehensive dataset containing over 400,000 records of insectivores and their associated microbes from 1903 to 2023. Our analyses showed that insectivores host a wide spectrum of 941 microbes, 60% of which are viruses and are predominantly found in the shrew and hedgehog families. Notably, human-associated viruses harbored by shrews and hedgehogs were phylogenetically closely related to those found in humans, suggesting potential bidirectional transmission between insectivores and humans. Moreover, virus-sharing networks revealed that insectivores held the second-most central position for virus sharing, just second to bats, among all mammalian orders. Insectivores had a high proportion of cross-order transmitted viruses, including many human-associated viruses. Dietary diversity, habitat diversity, and distributional traits (e.g. geographical range size, mean latitude, and urban adaptation status) emerged as the key ecological factors contributing to this cross-species virus transmission. Our findings highlight the microbial diversity present in insectivores, indicating this order may act as potential incubators for novel viruses capable of infecting mammals and spreading viruses of public health concern.

## Introduction

Most emerging infectious diseases (EIDs) in humans originate from pathogens hosted by non-human animals through zoonotic transmission^1,2^. Wild small mammals are the primary animal sources of these EIDs ^3–5^. Eulipotyphla (“true insectivores”, hereafter insectivores) is the third largest mammalian order, encompassing over 500 species across four taxonomic families: Erinaceidae (hedgehogs and moonrats), Soricidae (shrews), Solenodontidae (solenodons), and Talpidae (moles)^6,7^. Despite their small body size, insectivores can harbor a number of important zoonotic viruses, including but not limited to hantaviruses, Borna disease virus (BoDV), Middle East respiratory syndrome-related coronaviruses (MERS-related CoVs), severe fever with thrombocytopenia syndrome virus (SFTSV), and Langya henipavirus ^6–11^, which pose challenges to human, wildlife, and domestic animal health. However, the epidemiology and ecology of microbes in insectivores remain largely unexplored and poorly understood. This gap is a priority for the early detection and warning of EIDs.

Outside of zoonotic viruses, insectivores have been shown to carry a wide spectrum of microbes, with some species having even more viruses than well-known viral hosts, such as bats and rodents ^12–15^. This includes many genetically distinct viruses through reassortment or recombination ^16,17^. Some insectivore species (e.g., shrews) also have been shown to share many viruses with other wild species, even among host orders^1515^. Therefore, it is reasonable to speculate that insectivores may play a prominent role in the generation of novel pathogen variants, subsequently spreading them to other wild species and eventually posing a threat to public health through spillover and emergence.

Previous studies of insectivores have typically focused on specific host and microbe species in limited geographic areas, reducing their ability to generally assess the diversity and distribution of microbes in this order. A systematic investigation of microbes in insectivores is lacking. In addition, most studies have primarily aimed at discovering novel pathogens. Consequently, less is known the broader patterns of microbial diversity in insectivores and the factors that govern their transmission.

In this study, we integrated data from multiple sources to establish a comprehensive dataset of insectivores and their associated microbes. Our dataset comprised more than 400,000 records of insectivore hosts and 941 microbes from diverse habitats across the world, spanning from 1903 to 2023. This extensive dataset offered the unique opportunity to unveil the diversity and distribution of microbes in insectivores, including identifying viruses specifically with zoonotic potential. Moreover, we aimed to understand the role of insectivores in virus transmission across mammals more generally and to determine the ecological factors influencing this process.

## Methods

### Data collection

We collected data on Eulipotyphla, including information on the geographic locations of sampled insectivore hosts, the diversity and positivity (i.e., prevalence) of microbes, and genomic sequences of the microbes. These data were obtained from the literature, relevant websites, and the GenBank database (https://www.ncbi.nlm.nih.gov/genbank/). Electronic databases (PubMed and China National Knowledge Infrastructure Database) were searched by two independent reviewers (J.L. and L.H.) to extract data from research papers published before December 30, 2023, using a systematic protocol ^18^. Searches were conducted using both the Latin and common names (in English or Chinese) of each insectivore species. To avoid inconsistencies, specific Latin names were referenced according to the NCBI Taxonomy Database (https://www.ncbi.nlm.nih.gov/Taxonomy/Browser/wwwtax.cgi), disregarding the common names in the original publications. Publications were included if they contained relevant terms in the title, abstract, or keywords. Full-text publications were considered if they were in English or Chinese and provided detailed information on insectivores. Articles were excluded if they lacked sufficient information on the distribution and/or microbes of insectivores, or if they were duplicates. The detailed data extraction is shown in Fig. S1a. Geographic distributions of insectivores were extracted using the Global Biodiversity Information Facility ^19^ (https://www.gbif.org/) and mapped using ArcGIS (v10.8), supplemented by host geographic locations obtained from the literature. When exact locations were unavailable, the centroids of administrative areas were used. Additional microbial data of insectivores were retrieved from the Enhanced Infectious Disease Database (EID2)^20^ and GenBank database in April 2024. These microbes included viruses, bacteria, fungi, and protozoa. Genomic sequences of viruses in insectivores were obtained from GenBank for phylogenetic analyses. Relevant details were extracted, including insectivore species, location, virus classification, and the dates of sequence submission and release.

### Meta-analysis of microbe prevalence

We conducted meta-analysis to estimate the combined prevalence of positivity and 95% confidence intervals (CI) for microbes in insectivores. Studies reported the number of positive microbes using various assays, including PCR, next-generation sequencing, and microscopy. We calculated the pooled prevalence and 95% CI for microbe families/species in the insectivore order as well as per insectivore family and species. For microbes containing only one study, the prevalence was calculated without a 95% CI. At the microbe family level, if the total number of positive samples was available in the original publication, prevalence was calculated by dividing the total number of positives by the number of samples tested. If it was not possible to determine the total number of positive samples due to co-existence of multiple microbe species in individual samples, the highest prevalence for the microbe species was used to represent the entire family’s prevalence. If a study tested multiple insectivore species for microbes, results for each species were analyzed separately. We used the *I²* statistic to quantify heterogeneity in prevalence. If *I²* was less than 50%, a fixed-effects model was applied. Otherwise, a random effects model was used. Meta-analyses were conducted using the *meta* package (v7.0-0) in R (v 4.3.1).

### Transmission network and centrality measures

The mammal-virus and invertebrate-virus interactions were extracted from the EID2 and GenBank databases, including 34,253 interactions between 16,194 viruses and 3,731 host species. A bipartite network was constructed to connect each host order with its associated virus, comprising 18,497 interactions between 16,194 viruses and 141 host orders (20 mammalian orders and 121 invertebrate orders; Fig. S2). This bipartite network was then projected onto a weighted unipartite network, where nodes represented host orders connected by the number of shared viruses. The cross-order virus-sharing network included 723 host–host interactions between 1,039 viruses and 80 host orders (Fig. 4b). Additionally, networks were constructed at the host family, genus, and species level as well as for DNA viruses, RNA viruses, and human-associated viruses. Human-associated viruses were defined as those detected in humans ^21^.

Degree centrality, closeness centrality, eigenvector centrality, and the PageRank algorithm^22^ were calculated for each node in the viral transmission network. To adjust for host sampling bias, the number of PubMed citations with the Latin name of the host species and the term “virus” was used to correct the centrality metrics. The correction was based on the residuals obtained from regressing centrality on sampling effort ^23,24^. Specifically, centrality metric scores for each node were calculated using the number of shared viruses between two nodes as the edge weight, and these scores were then regressed against the sampling effort of each host. The residuals from each regression were used as the final centrality score for each node. The pairwise correlation coefficients between the four measures were all positive (Spearman’s *ρ* > 0.70, *P* < 0.05), indicating that the central hosts identified by different centrality indicators were similar (Fig. S3a). Therefore, closeness centrality was selected as the primary indicator for assessing central hosts in virus transmission.

### Ecological factors driving virus transmission

Life-history traits (body mass, longevity, litter size, and gestation time), foraging traits (habitat diversity and dietary diversity), distributional traits (range size and mean latitude), and urban adaptation status were collected from publicly available databases (Table S1-Table S3). Body mass and longevity data were sourced from the PHYLACINE ^25^ and Amniote databases ^26^, respectively. Data on litter size and gestation time were retrieved from Anage ^27^ and Amniote ^26^ databases. The size and mean latitude of hosts’ distribution range were obtained from PanTHERIA ^28^. Urban adaptation status for each species was determined using a published database on long-term urban adaptation in mammals^29^. Species classified as “dweller” or “visitor” were considered “urban-adapted”, while those not included in the urban-adapted dataset were coded as “non-urban”. Urban adaptation was thus treated as a binary variable at the host species level with species coded as 0 (non-urban) or 1 (urban). Body mass and range size were log-transformed to reduce skewness. These species-level trait values were used in the analysis of host species centrality, while the mean value (percentage for urban adaptation status) of a specific trait was calculated across the species within each genus or family to generate genus-or family-level trait values. Missing values were neglected when calculating genus-and family-level mean trait values.

Habitat use data were collected from the International Union for the Conservation of Nature (IUCN) database (http://www.iucnredlist.org). Dietary proportions were obtained from the PHYLACINE database. Habitat diversity at the species level referred to the number of habitats utilized by a species, whereas dietary diversity was calculated using the Shannon index based on dietary proportions. For both traits at the family and genus levels, dendrograms were generated for each trait, and Faith’s index (FD) was calculated by treating the species within the family or genus as a community. FD was calculated using the *picante* package^30^.

Phylogenetic trees for 75 host families, 198 host genera, and 664 host species of mammals were derived from the TimeTree database ^31^ and PHYLACINE databases. Based on these trees, we calculated the pairwise phylogenetic distances between species, genera, and families, which were included in the following analyses of host centrality. Phylogenetic signal for host centrality in virus transmission networks at the family, genus, and species level were calculated using Pagel’s λ from the *caper* package ^32–34^. A λ value of 0 indicates no phylogenetic signal, while λ of 1 suggests strong phylogenetic dependence.

Phylogenetic generalized least squares (PGLS) models were then applied to examine the effect of host traits on the centrality in viral sharing networks at the family, genus, and species levels. Candidate models were constructed following a full-as-possible model-building strategy, removing collinear predictor variables (i.e., Spearman’s ρ > 0.6) (Fig. S4). The number of predictor variables was limited to ensure that each estimated coefficient had at least 10 observations. Candidate models were compared using the Akaike information criterion (AIC), derived Akaike weights, and averaged coefficients when the ΔAIC of multiple models was less than two. PGLS models were fitted using the *phylolm* package, and model averaging was performed using the *MuMIn* package. Kruskal-Wallis tests were performed to compare median differences in host traits that affect host centrality.

### Phylogenetic analyses

For human-associated viruses found in insectivores, phylogenetic analyses were conducted based on the coding sequences (CDS) of conserved viral proteins, including the RdRp structural domain of RNA viruses, the capsid proteins of DNA viruses, and other specific genes of certain viruses (Table S4). Sequence alignment was performed using MAFFT (v7.525) ^35^. Maximum likelihood phylogenetic trees were constructed using the IQ-TREE (v2.2.6) ^36^, with branching support assessed using 1,000 SH-like approximate likelihood ratio test (SH-aLRT) replicates. Phylogenetic trees were visualized using *ggtree* (v4.3.1), *phangorn* (v4.3.1), and *ggplot2* (v4.3.1).

## Results

### Insectivore dataset

We established a multisource dataset comprising the geographic distribution of insectivores, their microbe diversity and prevalence, and the genomic sequences of these microbes from 1903 to 2023. The dataset included 432,941 location records of insectivores globally. Insectivores were documented in their natural habitats across 161 countries and territories on all continents except Antarctica (Fig. 1a). They were particularly recorded in Europe and North America, with high frequencies in the United Kingdom, France, the United States, Spain, and Germany. The distribution of records among insectivore host families varied considerably (Fig. S5). Shrews, hedgehogs, and moles had a similar number of records, while the solenodons accounted for a very small number of the total records (given only two extant species in this family, *Atopogale cubana* and *Solenodon paradoxus*).

**Figure 1.**
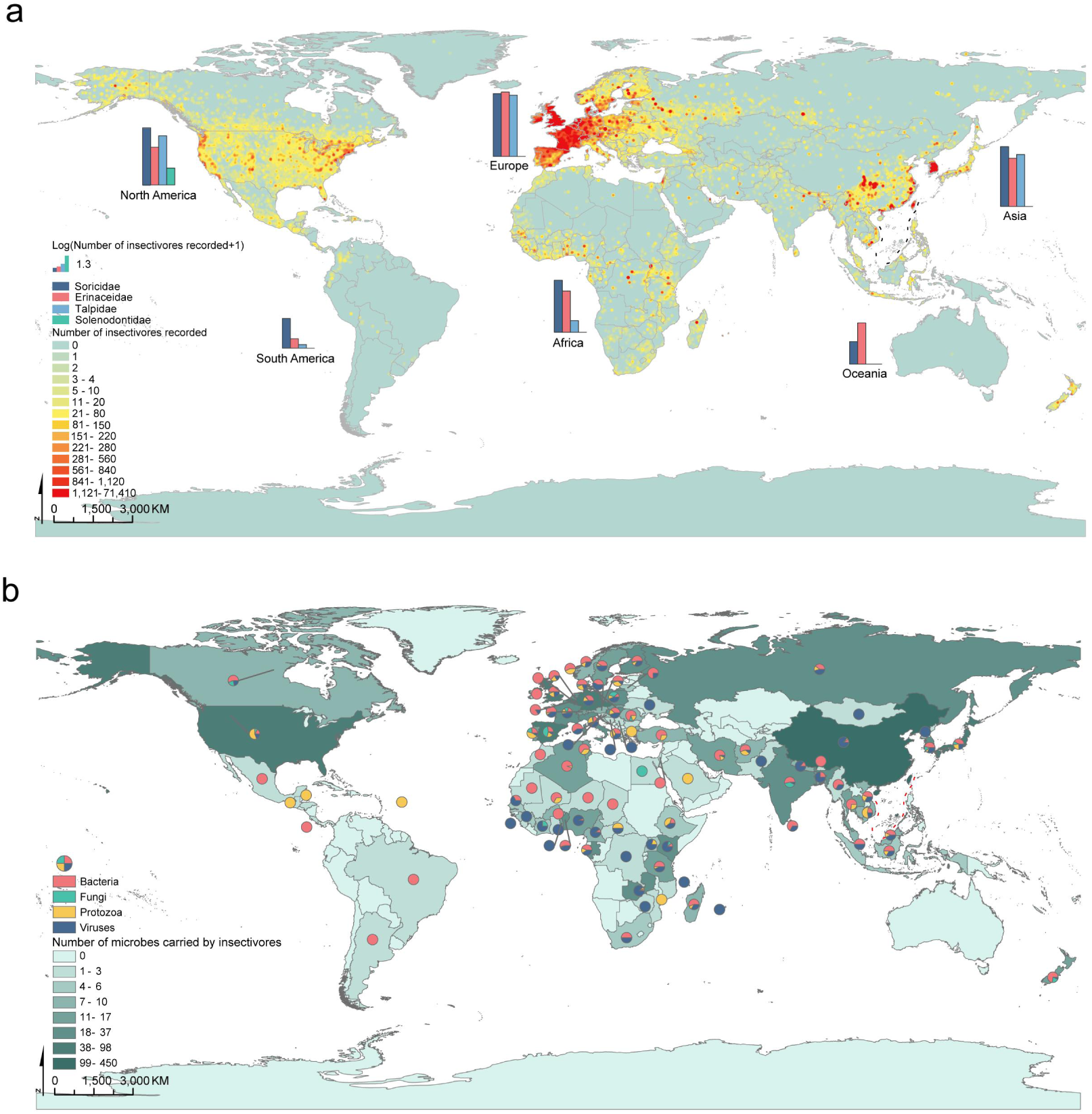
Geographical distribution of insectivores and their associated microbes. a) This map highlights the geographical distribution of insectivores, with darker shades indicating areas where higher numbers of insectivore species have been recorded. b) Geographic distribution of microbes harbored by insectivores. The accompanying pie chart displays the composition of four microbe categories (viruses, bacteria, protozoa, and fungi).

The dataset also contained 14,831 records and 10,245 genomic sequences of microbes in insectivores. The earliest microbial publication on insectivores was in 1905, and the number of related articles has gradually increased over time, reaching a total of 2,078 studies by 2023 (Fig. S1b). Notably, most genomic sequences of microbes in insectivores were obtained in the past two decades.

### Diversity and distribution of microbes in insectivores

We found a wide variety of microbes in insectivores, with an identified 941 unique microbes, including 569 viruses, 226 bacteria, 117 protozoa, and 29 fungi, across 94 orders and 173 families in 137 insectivore species (Fig. 2a, Fig. S6, and Table S5). Microbes from the families *Hantaviridae*, *Eimeriidae*, *Bartonellaceae*, and *Trypanosomatidae* were the most detected, each found in more than 25 insectivore species. Notably, 41.12% (234/569) of the viruses were invertebrate-specific.

**Figure 2.**
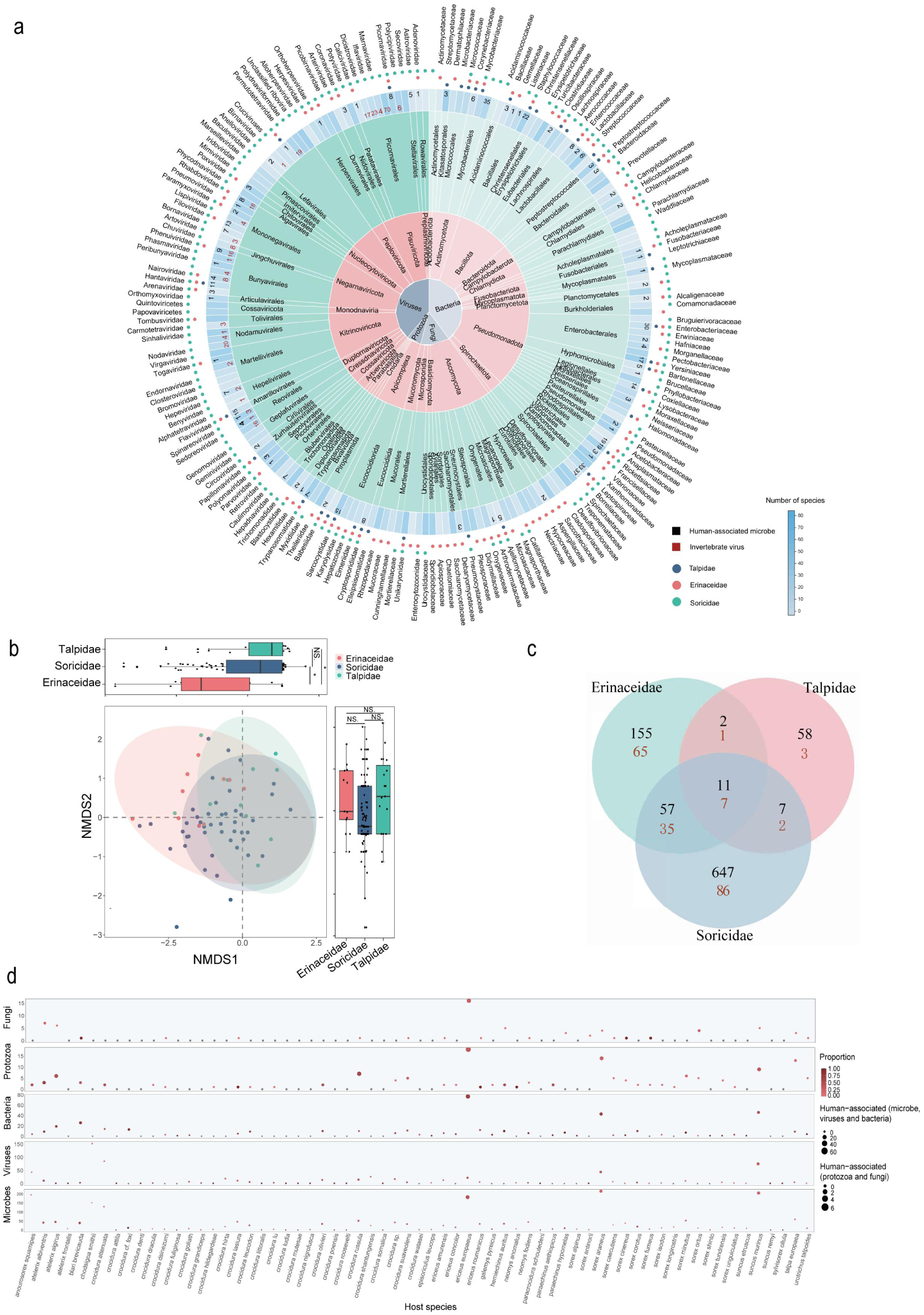
Overview of microbes harbored by insectivores. a) Classification of microbes in insectivores. The three inner circles represent the category, phylum, and order of the microbes, respectively. In the fourth circle’s grid, the number of species in the microbe family are represented by a gradient color. The number of human-associated microbe species and invertebrate virus species in each family are indicated by black and red number, respectively. The circle outside the fourth circle represents the host. The outermost names correspond to the microbe families. b) Non-metric multidimensional scaling analyses showing the variation in microbe composition among three host families within the insectivore order. Ellipses indicate 95% confidence intervals around the group centroid. c) Venn diagrams showing the number of microbes shared among three host families. Black and red denotes the number of microbes and the number of human-associated microbes, respectively. d) Profiles of microbes and human-associated microbes in insectivore species. The boxes indicate microbes, viruses, bacteria, protozoa, and fungi. The Y-axis represents the number of microbes harbored by the host. Circle size represents the number of human-associated microbes, and the circle color indicates the proportion of human-associated microbes within the microbes.

The diversity and composition of microbes varied substantially among the host families within insectivores (Fig. 2a-Fig. 2c). Shrews harbored the highest number of microbes, with 70% being viruses, predominantly from the families *Hantaviridae*, *Paramyxoviridae*, *Rhabdoviridae*, and *Phenuiviridae* (Fig. S6b). In contrast, hedgehogs had a higher proportion of bacteria, accounting for more than 50% of the microbes, mainly belonging to the families *Arthrodermataceae*, *Leptospiraceae*, *Enterobacteriaceae*, and *Borreliaceae* (Fig. S6c). Moles hosted fewer microbes, with the majority being protozoa from the family *Eimeriidae* (Fig. S6d). Shrews and hedgehogs tended to share more microbes, whereas only 11 microbes were common to all three host families (Fig. 2c). Certain shrew and hedgehog species had especially high microbial richness. The European hedgehog (*Erinaceus europaeus*), house shrews (*Suncus murinus*), and Eurasian shrew (*Sorex araneus*) each hosted over 100 microbial species (Fig. 2d). Some species were even capable of hosting a particularly high number of viruses. For example, Smith’s shrew (*Chodsigoa smithii*) hosted 152 viruses, 128 of which were invertebrate-specific viruses (Fig. 2a).

Microbes in insectivores were documented on six continents, with a predominant presence in Asia, Europe, and Africa (Fig. 2b). Microbial composition varied by continent, with viruses and bacteria mainly recorded in Asia, Europe, and Africa. Fungi and protozoa were mainly found in Europe and East Asia. The geographic distribution of microbes also varied across different host families (Fig. S7).

### Microbial prevalence

We next sought to determine the prevalence of microbes in insectivores, revealing considerable differences across microbe families (Fig. S8). Among the virus families, the *Circoviridae* (91.30%, n=1 study) and *Arteriviridae* (90.79%; 95% CI: 65.03-100%) had higher prevalence. Microbial prevalence varied significantly among different host families as well, with higher prevalence in hedgehogs (generalized linear model [GLM], β = 1.24, *P* < 0.001) and moles (β = 1.25, *P* < 0.001; Fig. 3 and Fig. S9). Viruses in shrews mainly belonged to the *Hantaviridae*, *Flaviviridae*, and *Paramyxoviridae* families, with prevalence of specific zoonotic viruses being 46.67% (Xinyi virus, n=1 study), 33.33% (*Orthoflavivirus powassanense*, n=1 study), and 47.57% (*Jeilongvirus apodeme*, 95% CI: 0-100%). In hedgehogs, high-prevalence viruses included *Betacoronavirus erinaceus* (54.35%; 95% CI: 26.02-82.67%), Mecsek mountains virus (45%, n=1 study), and *African pygmy hedgehog adenovirus 1* (43.89%; 95% CI: 0-100%). By contrast, viruses in moles were almost exclusively from the *Hantaviridae* family, including Academ virus (77.78% prevalence, n=1 study) and *Orthohantavirus asamaense* (60% prevalence, n=1 study). Most bacterial and fungal species were present only in hedgehogs (Fig. 3). *Isospora rastegaievae* had 80% prevalence (n=1 study), *Anaplasma phagocytophilum* had 71.31% prevalence (95% CI: 38.83-100%), and *Bacteroides fragilis* had 64.17% prevalence (95% CI: 0-100%). Protozoa with prevalence greater than 50% were almost exclusively present in moles (Fig. 3).

**Figure 3.**
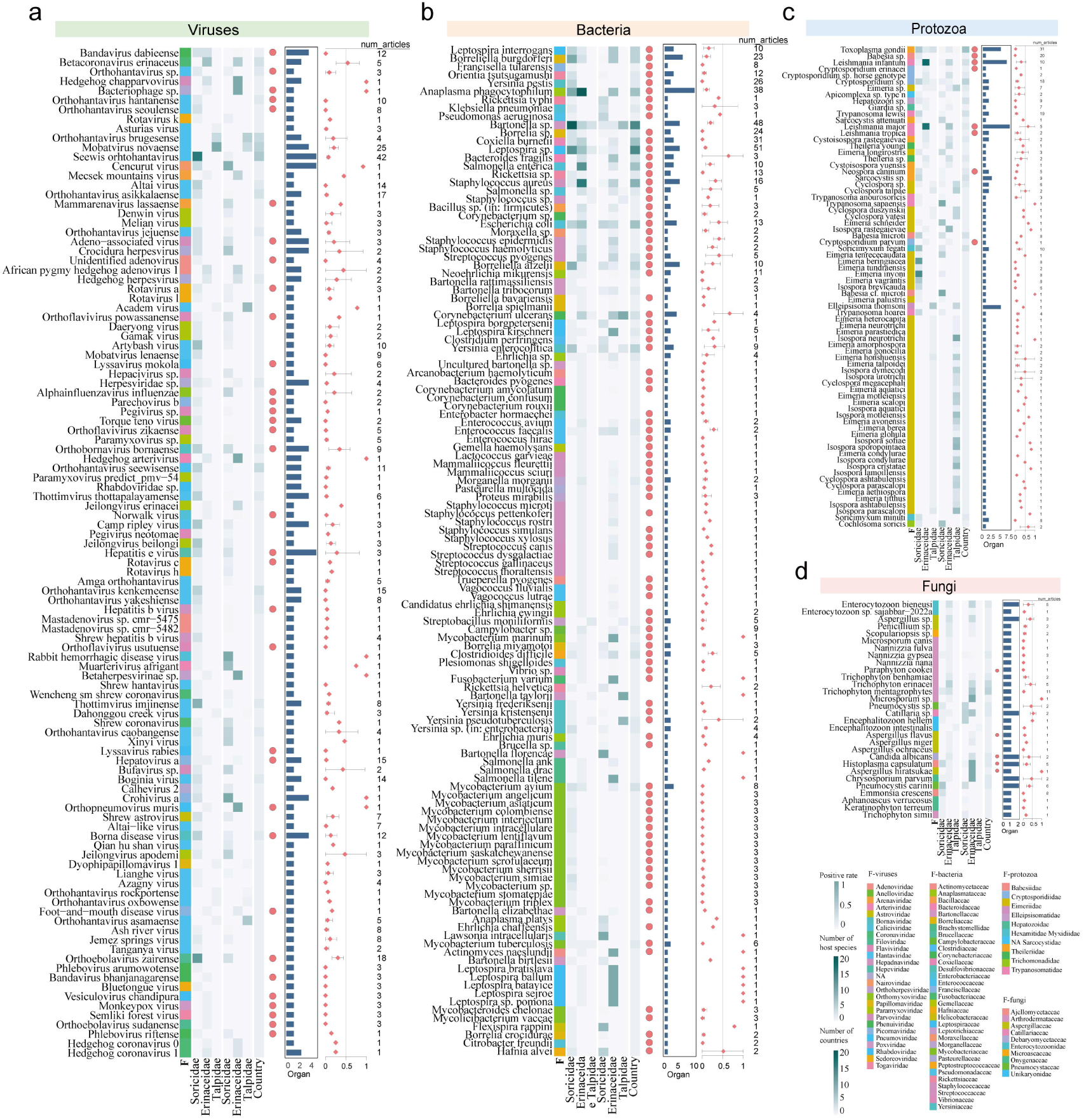
Prevalence of microbes harbored by insectivores. a) viruses, b) bacteria, c) protozoa, d) fungi. In the heat map, the three columns on the left represent the number of host species within each host families that carry the microbe, the three middle columns represent the prevalence of the microbe from the host families after meta-analysis, and the column on the right represents the number of countries where the microbe has been detected. The red dots on the right of the heatmap indicate human-associated microbes. The histograms denote the number of organs in which each microbe has been detected. The forest plots illustrate the prevalence and the 95% confidence intervals for microbes in insectivores after meta-analysis.

We also investigated the distribution of microbe prevalence across organs and continents (Fig. 3 and Fig. S10). Prevalence showed considerable variation in organ distribution, with significantly lower prevalence in lung (GLM, β =-1.11, *P* < 0.001), liver (β =-1.35, *P* < 0.001), and spleen (β =-0.95, *P* = 0.01) than gut. *Hantaviridae* viruses were predominantly detected in the lungs, while microbial species from the *Anaplasmataceae* and *Leptospiraceae* were found at high prevalence in spleen and kidney samples, respectively (Fig. S10). Moreover, we observed substantial differences in prevalence of multiple microbes in different organs within the same host. Cencurut virus and hepatitis E virus in the house shrew (*Suncus murinus*), and *Orthobornavirus bornaense* and BoDV in the bicolored shrew (*Crocidura leucodon)*, were present in two or more organs. Notably, there were several microbes exhibiting multi-organ distribution across species and countries. *Seewis orhtohantavirus* was detected in four organs in 16 shrew species from six European countries, with a prevalence of 7.15% (95% CI: 3.40-10.90%). *Toxoplasma gondii* was found in five organs in six shrew species, two hedgehog species, and three mole species, with a prevalence of 2.59% (95% CI: 1.27-3.92%; Fig. 3). Additionally, *Anaplasma phagocytophilum* and *Staphylococcus aureus* were detected in multiple organs and were widespread across continents (Fig. 3 and Fig. S10).

### Viral sharing networks

To better characterize the role of insectivores in virus transmission, we constructed host–virus correlation and virus–sharing networks. Insectivores harbored an average of 202 viruses per family, which was lower compared to ticks (Ixodida; 734), bats (Chiroptera; 366.36), and primates (Primates; 326.15). Meanwhile, the average number of viruses per genus in insectivores was 47.71, ranking second only to odd-toed ungulates (Perissodactyla; 86.0), primates (85.60), and bats (62.88) among mammals (Fig. 4a). Topological analyses of the virus-sharing network revealed that insectivores held the second-most central positions in the network, following bats among all mammalian host orders (Fig. 4b and Fig. 4d). However, sampling effort (i.e. citation counts) for insectivores was substantially lower than other mammals, representing only 3% of that for primates, rodents (Rodentia), bats, and carnivores (Carnivora; Fig. S3b).

**Figure 4.**
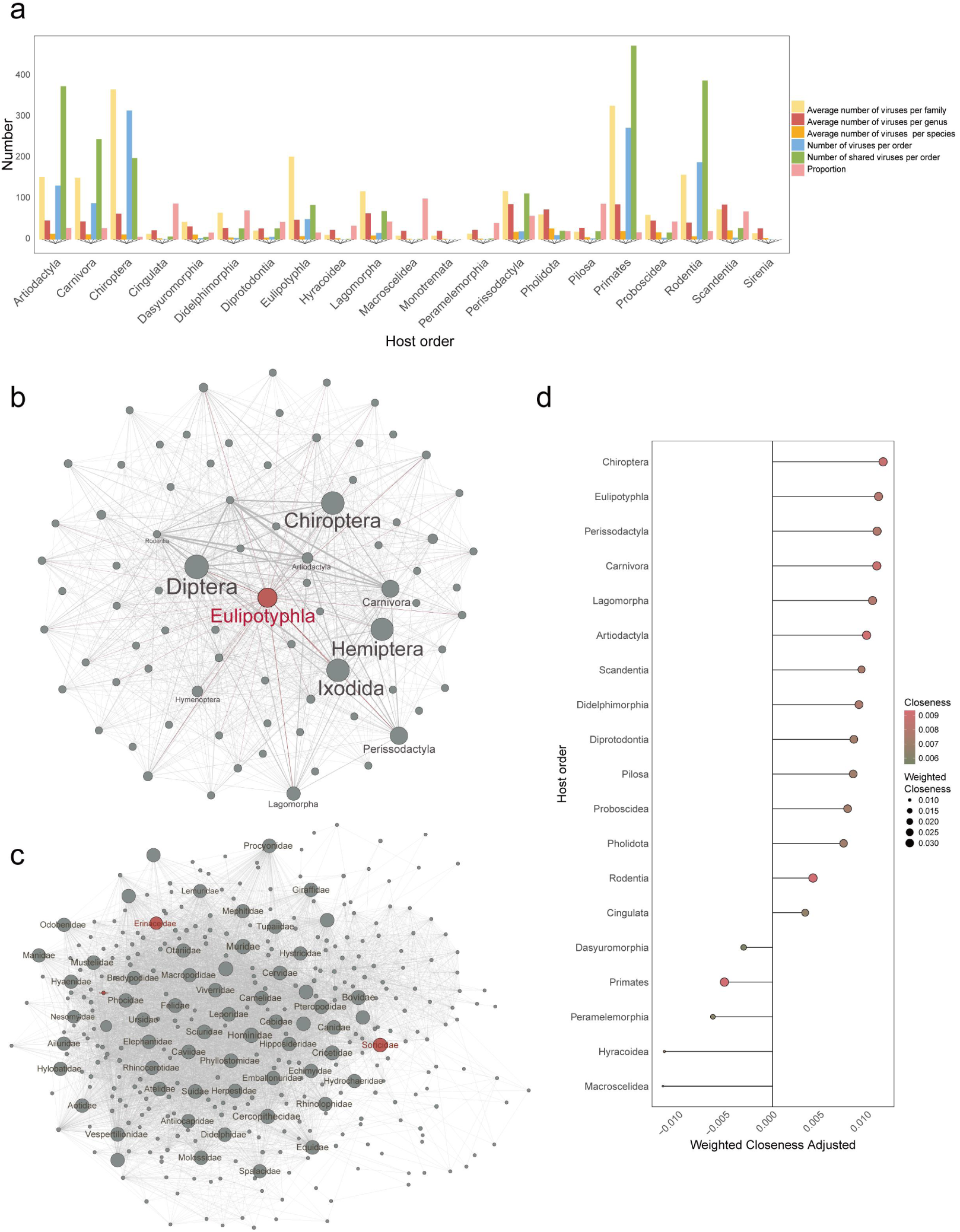
Transmission of viruses among mammals and invertebrates. a) Number of viruses harbored and shared by each mammalian order. “Number of viruses per order” indicates the number of viruses in the host order divided by 10. “Proportion” represents the proportion of shared viruses to the total number of viruses. b) Cross-order virus sharing network. c) Cross-family virus sharing network. The size of each node depicts the closeness centrality. d) Closeness centrality of mammalian orders in the cross-order virus sharing network. The x-axis represents the weighted closeness centrality adjusted. The circle size represents the weighted closeness centrality. The circle color represents the closeness centrality.

Virus transmission likely occurred between insectivores and other hosts at multiple taxonomic levels: species (122 viruses), genus (108 viruses), family (86 viruses), and even order (84 viruses). We found that insectivores had a higher proportion of cross-order viruses (16.94%) than bats (5.36%) and true bugs (Hemiptera; 16.45%), comparable to that of primates (17.37%; Fig. 4a). The 84 cross-order transmitted viruses were shared with hosts from 34 distinct orders, such as rodents, primates, and even-toed ungulates (Artiodactyla). The number of shared viruses between insectivores and other orders was significantly correlated with the phylogenetic distance of the hosts (Spearman’s *ρ* = 0.555, *P* = 0.002). Notably, in insectivores, human-associated viruses accounted for more than half (43/84, 51.19%) of the cross-order transmitted viruses, although they represented only a small portion (43/496, 8.67%) of the total viruses. Insectivores were more likely to share RNA viruses rather than DNA viruses: of the 84 cross-order transmitted viruses, 72 were RNA viruses and 12 were DNA viruses. In the RNA virus-sharing network, insectivores held the second-most central positions after bats among all mammalian orders, while they ranked fifth in the DNA virus network (Fig. S3c and Fig. S3d).

Viruses in insectivores were mainly found in the shrew and hedgehog families, prompting further investigation into virus-sharing patterns at different host taxonomic levels. The virus-sharing network at the host family level revealed that shrews and hedgehogs were more likely to transmit viruses to other hosts, ranking in the top 7% and 18% of the host families, respectively. In contrast, moles ranked in the top 53% (Fig. 4c). At the host species level, the house shrew, Eurasian shrew, and European hedgehog ranked in the top 12% of the 2,942 included species. We observed a similar pattern when only considering human-associated viruses (Fig. S3). These findings suggest that insectivores, particularly the shrew and hedgehog families, contributed significantly to virus transmission.

### Ecological factors affect virus transmission

To further identify the ecological drivers governing the centrality of insectivores in virus transmission networks across families, genera, and species, we conducted phylogenetic comparative analyses using PGLS models. The centrality of host families (λ= 0.533) and species (λ= 0.265) exhibited moderate and weak phylogenetic signals, respectively, whereas no phylogenetic signal was detected at the host genus level (λ= 0). We identified nine, 12, and 13 competitive PGLS models (ΔAIC < 2) at the family, genus, and species levels, respectively (Table S6-Table S8). Longevity was the most significant factor influencing host centrality at the family level (β = 0.53, *P* < 0.001), ranking second at the genus level (β = 0.16, *P* = 0.049) and third at the species level (β = 0.16, *P* = 0.004; Fig. 5a-Fig. 5c); these results indicate that longevity may be a common trait among central hosts in virus transmission. Habitat diversity was also positively associated with host centrality at both the genus and species levels (Fig. 5b and Fig. 5c). In addition, host centrality at the family level was positively correlated with dietary diversity and negatively correlated with phylogenetic distance (Fig. 5a). At the species level, higher host centrality was correlated with larger geographic range size, higher mean latitude, and urban-adapted status (Fig. 5c).

**Figure 5.**
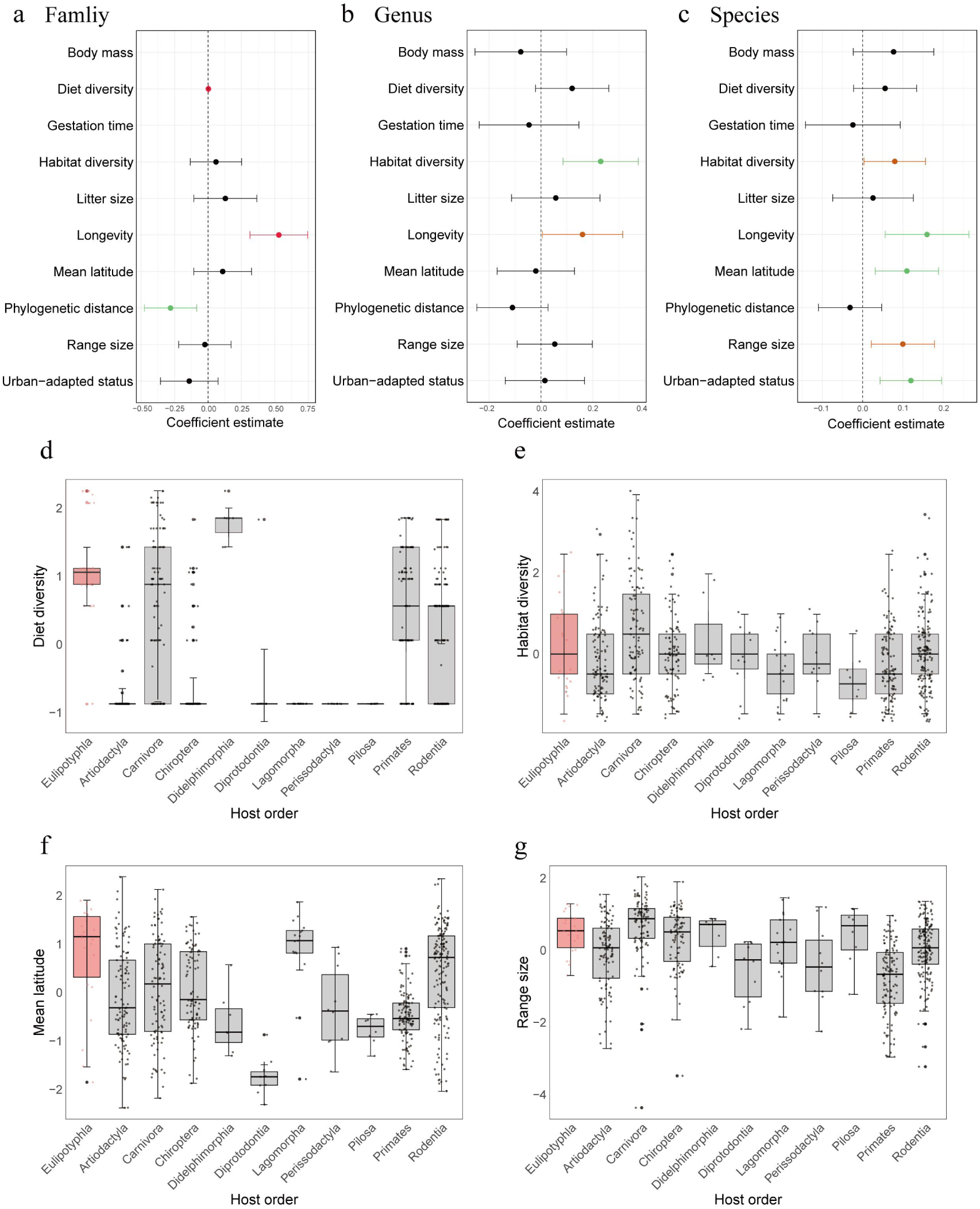
Summary of the ecological traits that affect virus sharing among mammals at different taxonomic levels. The coefficients and corresponding *p*-values were estimated using the best PGLS models at host a) family level, b) genus level, and c) species level. The circles indicate the estimates and lines show 95% CIs. Red: *P*-values < 0.001, green: *P*-values < 0.01, dark orange: *P*-values < 0.05, and black: *P*-values > 0.05. d) Diet diversity, e) habitat diversity, f) mean latitude, and g) range size of host species from different mammalian orders.

We also examined differences in host traits across orders to broadly assess distinct ecological features of insectivore species (Kruskal-Wallis tests: *P* < 0.05). Insectivores have the second highest dietary diversity among the 11 mammalian orders, following marsupials (Didelphimorphia; Fig. 5d). Although the median habitat diversity of insectivores is in the middle range (Fig. 5e), moles and shrews rank in the top 3.95% and 21.05%, respectively. In addition, the mean latitude of insectivores is higher than that of all other mammalian orders (Fig. 5g), whereas their geographic range size is smaller than only carnivores, marsupials, and sloths and anteaters (Pilosa; Fig. 5f).

### Shrews and hedgehogs carry human-associated viruses closely related to those in humans

Phylogenetic trees revealed that 11 human-associated viruses found in shrews and hedgehogs were closely related to those identified in humans. These viruses can be divided into two major categories based on their transmission modes (Fig. 6).

**Figure 6.**
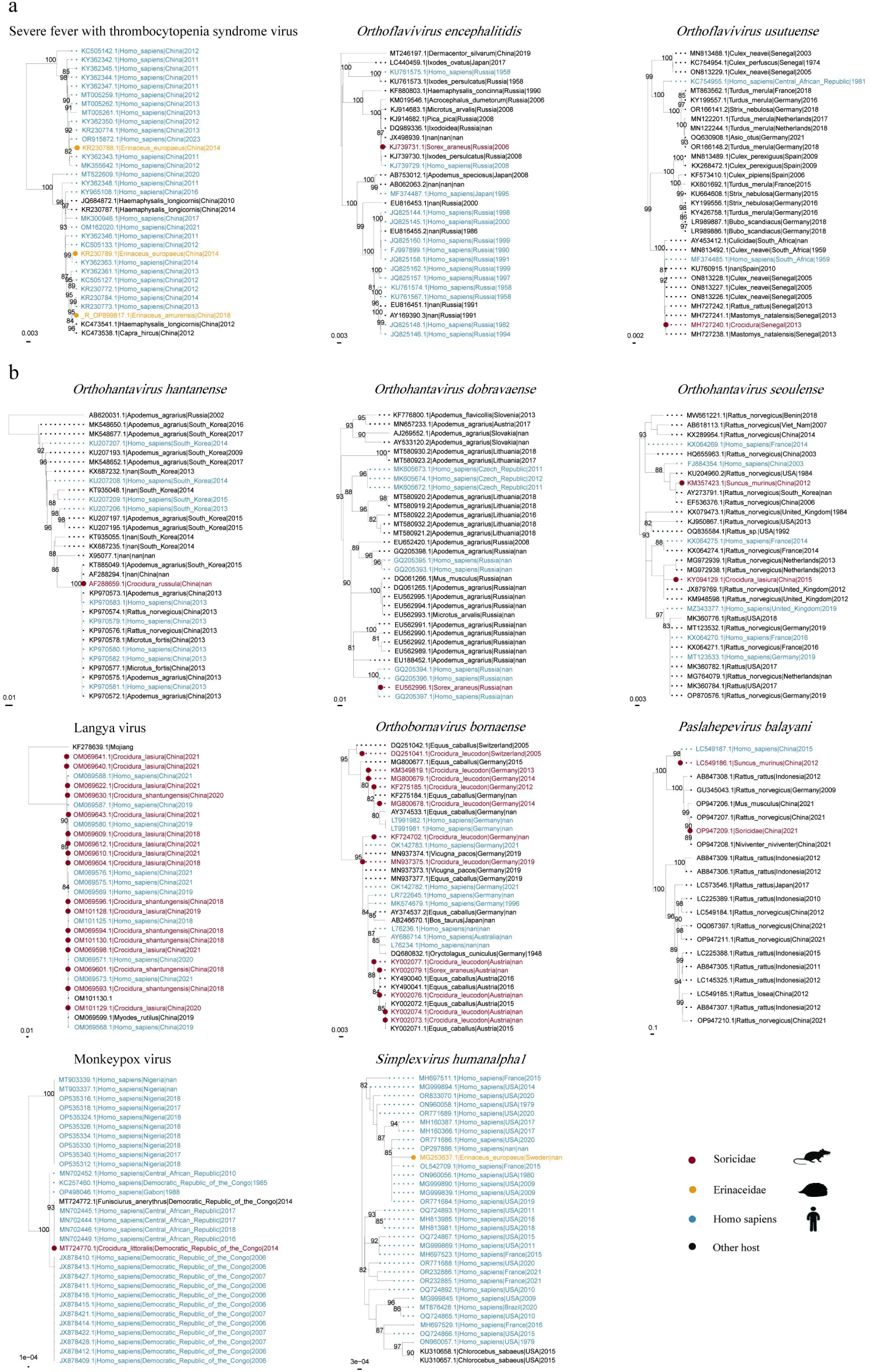
Maximum likelihood phylogenies of human-associated viruses harbored by insectivores. a) Phylogenies of vector-borne viruses. b) Phylogenies of other viruses. The host species for each virus is represented by different colors as shown in the key. Bootstrap support values are displayed at the node. The length of the branches in the tree are proportional to the number of nucleotide substitutions per site as indicated by the scale.

The first category consisted of three vector-borne viruses likely transmitted to humans from insectivores via ticks (Fig. 6a). SFTSV in European hedgehogs clustered with strains from humans and *Haemaphysalis longicornis* in China (nucleotide identity 94.94-99.85%). Hedgehogs may serve as amplifying hosts of SFTSV, with *Haemaphysalis longicornis* as the primary transmission vector. In addition, *Orthoflavivirus encephalitidis* in Eurasian shrews and *Orthoflavivirus usutuense* in white-toothed shrews (*Crocidura*), both belonging to the *Flaviviridae* family, were phylogenetically related to human strains with nucleotide identities of 99% and 97.30-99.85%, respectively. *Orthoflavivirus encephalitidis*, the causative agent of tick-borne encephalitis, is endemic in Eurasia. The virus in shrews was also closely related to those from ticks, primarily transmitted by *Ixodes ricinus* and *Dermacentor reticulatus*.

The second category comprised eight non-vector-borne viruses, showing potential bidirectional transmission between insectivores and humans (Fig. 6b). The newly identified Langya virus in shrews was closely related to those in febrile patients (nucleotide identity 99.22-100%); shrews are considered the potential natural reservoirs for this virus^37,38^. *Orthohantavirus hantanense*, *Orthohantavirus seoulense*, and *Orthohantavirus dobravaense*, all within the *Hantaviridae* family, were also closely related to human strains (nucleotide identity 94.57-99.61%, 90.50-99.50%, and 95.40-97.93%, respectively). Similar relationships were observed for *Orthobornavirus bornaense*, *Paslahepevirus balayani*, and monkeypox virus. Notably, our phylogeny revealed that *Simplexvirus humanalpha1* in the European hedgehog clustered within a group of human strains (nucleotide identity 99.05-99.87%), suggesting potential spillback to European hedgehogs, as humans are the primary hosts for this virus^39^.

## Discussion

In this study, we integrated data from multiple sources to reveal the distribution and diversity of microbes in insectivores and explored their role in virus transmission.

### Role of insectivores as viral hosts

We observed that insectivores, particularly shrews, harbor a wide spectrum of viruses. Some shrew species may even carry a greater diversity of viruses compared to bats and rodents. Growing evidence suggests that insectivores serve as natural reservoir hosts for multiple viruses ^40–42^. Although relatively little is known about which biological traits of insectivores may facilitate this high viral diversity, it is likely that they share certain characteristics with both bats and rodents that may allow for the efficient maintenance of viruses. Like rodents, insectivores have a fast life history, which may enhance transmission through population dynamics ^43^, but, like bats, they also have high metabolism, exhibit torpor (as do some rodents), and may have unique immune responses to viral infection. Such bat-like factors likely contribute to the persistence of chronic infection or decrease viral replication through within-host dynamics^5,44^. Historically, our understanding of virus diversity in wild small mammals has been strongly skewed towards bats and rodents, with insectivores representing only a small fraction of mammals studied ^45–47^. This can only in part be explained by the greater species richness of these two orders ^48^, given the likewise speciose nature of the Eulipotyphla. Comprehensive investigation of the virome in insectivores, representing multiple species, distinct ecological habitats, and at the scale of individual animal, could offer a more complete view of the virus diversity in this order and offer insights into the emergence of zoonotic infections.

### Potential incubator for novel viruses that could infect mammals

Our study reveals that insectivores carry a large proportion of invertebrate-associated viruses. This is likely due to the diet of species in this order, many of which have invertebrates as the majority of their diet. Moreover, a high abundance of arthropod-specific or related viruses has been found in various organs in shrews ^15^, suggesting true infection rather than only colonization of the viruses. The arthropod-specific or related viruses may have evolved within insectivores and expanded their host range to other mammals.

Moreover, the diversity of viruses found in insectivores creates favorable conditions for viral genome recombination and reconfiguration. Previous studies have demonstrated the presence of viral recombination or reassortment in shrews ^16,17,49^, including reassortment between divergent virus strains ^50^. Invertebrate-associated viruses could potentially recombine or reassort with mammalian viruses in shrews, leading to the emergence of novel viruses or variants. A comparable example has been observed in swine, which have been hypothesized to serve as “mixing vessels” for the evolution of influenza A viruses ^51^. Swine are susceptible to infection with both avian and human influenza viruses, enabling the generation of genetically novel and pandemic strains through reassortment. For instance, the 1957 and 1968 pandemic strains influenza A viruses likely arose through reassortment between an avian influenza strain and the strain circulating in humans at the time ^52–54^. Given this, it is plausible to suspect that shrews might similarly act as potential incubators facilitating the reconfiguration of viral genomes between mammalian and non-mammalian viruses. Therefore, continuous monitoring and characterization of viruses circulating in insectivores are crucial for providing early warnings about the emergence of novel viruses or variants with increased pathogenicity or zoonotic potential.

### Role of insectivores in cross-species transmission among mammals

Consistent with previous studies ^55,56^, we found that bats, ungulates, and carnivores play important roles in the transmission of viruses among mammals. However, our study differs from previous studies in providing new insights into virus transmission across mammals. We found that insectivores, specifically shrews and hedgehogs, contributed significantly to virus-sharing networks, in particular by hosting many cross-order transmitted viruses.

We next explored the factors contributing to the central role of insectivores in viral-sharing networks by investigating the drivers at different taxonomic levels (i.e. family-, genus-, and species-level networks). We found that habitat diversity was positively correlated with hosts’ centrality at both genus and species levels. For insectivores, their global distributions in diverse ecological habitats allow them to be exposed to more viruses and thus enhance their potential to spread viruses across different environments. Dietary diversity was also positively associated with hosts’ centrality at the family level. On average, invertebrates account for about two-third of insectivores’ diet. The diverse diet of insectivores could facilitate virus transmission by consuming a wide range of invertebrates, which is consistent with the identification of many invertebrate-associated viruses. On the other hand, carnivores consuming virus-infected insectivores would also create opportunities for cross-species transmission. These predators could potentially act as amplifying hosts, contributing to the persistence and wider dissemination of these viruses. In addition, our data showed that urban-adapted status can facilitate virus transmission at the species level, which is consistent with previous studies ^33,57^. Many shrew and hedgehog species are adapted to urban environments.

Previous studies have shown that viruses are more likely to spread between phylogenetically related species ^56,58–61^. We likewise found a negative relationship between phylogenetic distance and frequency of viral sharing among mammal species, but this relationship was only detected at the family level. Insectivores are ancient mammals, and viruses that evolved in insectivores may use conserved cellular receptors that enhance their ability to transmit viruses to other mammalian orders ^62^. Finally, we found that host longevity was positively correlated with network centrality at all three levels. This might be explained by long lifespans facilitating greater cumulative exposure to viruses.

### Carrying viruses of public health concern

Lastly, our phylogenetic analyses revealed that certain viruses found in shrews and hedgehogs, such as Langya henipavirus, SFTSV, and BoDV-1, were closely related to viruses infecting humans. These viruses have been documented to transmit via various transmission routes and are known to cause diseases in humans ^8^. For example, hantaviruses can be transmitted to humans through inhalation of respiratory secretions, leading to serious diseases such as haemorrhagic fever with renal syndrome and hantavirus pulmonary syndrome. Hedgehogs, often heavily infested with ticks, harbor multiple tick-borne viruses ^14,63^. They have been recognized as amplifying hosts of SFTSV^64,65^, which can cause a wide range of clinical outcomes and has a case-fatality rate of 12–50% ^66–68^. Shrews have been identified as natural hosts of the newly discovered Langya henipavirus, which is closely related to the two well-known henipaviruses, Hendra virus and Nipah virus ^8^. Langya henipavirus can cause respiratory symptoms and has been associated with febrile illness in humans, particularly in farmers who reported contact with shrews.

In addition, we observed that several human-associated viruses, such as SFTSV, exhibited high prevalence and multi-organ distribution in insectivores. Such viruses showed a broad host range and possess a high probability of crossing species barriers. Given the wide geographical distribution of insectivores, along with the high prevalence, multi-organ distribution, and various transmission modes of their viruses, there is an elevated likelihood of further zoonotic spillover events. Given these risks, we recommend surveillance of high-risk human populations, such as agricultural workers who may come into regular contact with shrews and hedgehogs, to assess the zoonotic potential of these viruses.

Furthermore, shrews and hedgehogs are not restricted to rural areas. They can thrive near humans in urban settings and thus could carry and sustain the natural circulation of their zoonotic viruses within suburban and urban environments ^69,70^. Human activities that increase exposure to urban-adapted shrews heighten the risk of virus spillover to humans or domestic animals. Moreover, shrews tend to display bolder behaviours in urban settings ^71^, which may in turn increase their contact with humans and increase the risk of virus spillover. Pathogen surveillance at the wildlife-human interface in urban settings should be considered as a critical component of public health interventions aimed at controlling infectious diseases.

## Conclusions

In summary, our findings demonstrate that insectivores harbor a wide spectrum of microbes, particularly viruses. Shrews and hedgehogs in particular may serve as potential incubators for novel viruses capable of infecting mammals, and these families play a key role in the spread of viruses that pose significant public health risks. The repeated emergence of pandemic viruses that have originated from wild animals underscores critical gaps in current biosecurity systems. Subsequently, global surveillance and further research on insectivores should be intensified to better assess the risks associated with zoonotic transmission and to guide more effective strategies for disease prevention and control.

## Acknowledgements

We gratefully acknowledge Professor Xuejie Yu and Xuelong Jiang for helpful discussion on this study. This study was supported by the National Natural Science Foundation of China (32470561, 32271605), Taishan Scholars Project (tsqn202306003), and Shandong Excellent Young Scientists Fund Program (2022HWYQ-056). DJB was supported by the National Science Foundation (BII 2213854).

